# ReDirection: a numerically robust R-package to characterize every reaction of a user-defined biochemical network with the probable dissociation constant

**DOI:** 10.1101/2023.07.12.548670

**Authors:** Siddhartha Kundu

## Abstract

Biochemical networks integrate enzyme-mediated substrate conversions with non-enzymatic complex formation and disassembly to accomplish complex biochemical and physiological function. The multitude of theoretical studies utilizing empirical/clinical data notwithstanding, the parameters used in these analyses whilst being theoretically sound are likely to be of limited biomedical relevance. There is need for a computational tool which can ascribe functionality to and generate potentially testable hypotheses for a biochemical network. “ReDirection” characterizes every reaction of a user-defined biochemical network with the probable dissociation constant and does so by combinatorially summing all non-redundant and non-trivial vectors of a null space generated subspace from the stoichiometry number matrix of the modelled biochemical network. This is followed by defining and populating a reaction-specific sequence vector with numerical values drawn from each row of this subspace, computing several descriptors and partitioning selected terms into distinct subsets in accordance with the expected outcomes (forward, reverse, equivalent) for a reaction. “ReDirection” computes the sums of all the terms that comprise each outcome-specific subset, maps these to strictly positive real numbers and bins the same to a reaction-specific outcome vector. The p1-norm of this vector is the probable dissociation constant for a reaction and is used to assign and annotate the reaction. “ReDirection” iterates these steps recursively until every reaction of the modelled biochemical network has been assigned an unambiguous outcome. “ReDirection” works on first principles, does not discriminate between enzymatic and non-enzymatic reactions, offers a mathematically rigorous and biochemically relevant environment to explore user-defined biochemical networks under naive and perturbed conditions and can be used to address empirically intractable biochemical problems. The utility and relevance of “ReDirection” is highlighted with an investigation of a constrained biochemical network of human Galactose metabolism. “ReDirection” is freely available and accessible from the comprehensive R archive network (CRAN) with the URL (https://cran.r-project.org/package=ReDirection).

## 1 Introduction

An undirected biochemical network is converted into a pathway by a combination of physicochemical (temperature, pH, compartmentalization)- and biochemical (small molecule effectors, shared intermediates, feedback)-factors. Despite the availability and accessibility of advanced data analytical tools, true mechanistic insights into the manner in which a biochemical network accomplishes complex function is unclear [1-5]. An essential first step in the analysis of a biochemical network is the construction of a suitable model. This is usually data-driven and coarse-grained where nodes can represent proteins, genes or cells and edges indicate lines of supporting evidence (empirical, “omics”-datasets, co-expression data, text mining, knowledge-based databases) [1-10]. Analysing such a network will result in several network-specific characteristics such as the clustering coefficient and path distance [6-10]. This initial characterization is complemented by a library of equally plausible outcomes all of which are made to approximate the original architecture [10, 11]. Inverse modelling, for a dataset, will generate several possible candidate causal network models, allow hypothesis testing and may potentially be more informative [6-12].

Causal networks (CNs) are probability-based and can model alternate scenarios for every node of a small network whilst concomitantly ascribing specific states to each node [12]. Although CNs have had considerable success in investigating real world problems, inferring biochemical function from a network of genes/proteins/metabolites remains challenging [12]. For example, a causal network is usually modelled as an “acyclic”-graph which is in complete contrast to the plethora of feedback (positive, negative) mechanisms and reverse reactions that exemplify biochemical systems [12]. CNs are also inferential, modelled as a homogenous Poisson’s process (discrete event, discrete domain) and inherently Markovian [10, 12]. Biochemical function, on the other hand is dependent on thresholds (signal transduction, pattern receptors), characterized by minor perturbations and is memory-driven, all of which are better modelled as continuous events or variables in discrete time. CNs, to be truly informative also require a significant amount of data, *a priori*; a major limitation of CN usage in modelling biochemical networks. These arguments notwithstanding, CNs have contributed to well defined observables in the presence of ample empirical data such as phenotype mapping along with dose- and stimulus-driven response of genes [13-17]. CNs of genes and proteins result in lists which can be utilized for large scale data mining (parameter selection, candidate genes) and/or analytics as in precision medicine and biomarker profiling [2, 3, 13-17].

Unlike data-driven modelling, optimization- and enumeration-based strategies can be used to investigate and characterize a biochemical network from first principles and at near steady-state [18-26]. Flux-based algorithms (flux balance analysis, flux-variable analysis, regulatory on-off minimization, minimization of metabolic adjustment), will maximize or minimize the biomass of a metabolite of interest and can be used to investigate the effects of deletions and other perturbations on the flux of metabolites through a large network [18-22]. The numerical enumeration of elementary flux modes- and vectors-, along with extreme pathway analysis can be used to derive meaningful information such as “metabolic”-hubs and smaller subsets of cooperating reactions from biochemical networks [23-26]. The integration of empirical data (“omics”-studies, spectroscopy, pulse-chase experiments) into theoretical studies is referred to metabolic flux analysis (MFA) and is utilized in biotechnological applications to regulate the biomass of a preferred reactant/product [27-29].

The aforementioned limitations of data-driven models or biomass optimization advocates the need for a computational tool which is unbiased, biochemically relevant, theoretically sound and numerically robust and generate testable hypotheses. The dissociation constant is a non-negative real number which can be mapped to several biochemically relevant outcomes of a reaction (forward, reverse, equivalent, tight binding) [30-35]. The probable dissociation constant, on the other hand is a numerical measure that can be computed from the null space generated subspace of a stoichiometry number matrix of a biochemical network and possesses several desirable properties of the true dissociation constant [36]. “ReDirection” is an R-package which will compute the probable dissociation constant for every reaction of a user-defined biochemical network. The manuscript introduces some of the principles and definitions that will be used by “ReDirection” to characterize the modelled biochemical network. An outline of the functions used by “ReDirection”, their dependencies, rationale and usage is presented. A stepwise description and brief analysis of the algorithm that “ReDirection” deploys is also described. This will be followed by numerical studies on a constrained biochemical network of human Galactose metabolism. The manuscript concludes with a summary of the salient features, limitations and possible future studies which may utilize “ReDirection”.

## 2 Methods

### 2.1 Definitions, concepts and notation relevant to comprehending the functionality of “ReDirection”

The algorithm that is deployed by “ReDirection” is mathematically rigorous, biochemically relevant and has been extensively discussed [36]. Briefly, a biochemical network is modelled as an instance (𝓅_z_) of *i*-indexed (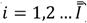) *r*-reactions that results from selected combinations of *j*-indexed (*j* = 1,2 … *J*) *m*-reactants and is the sparse stoichiometry number matrix (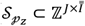) (**Defs. (1a-1d)**) [36]. The modelled biochemical network is constrained with pre-defined and dynamically determined or network-specific criteria (**Def. (2)**) [36]. The former includes lower bounds for the numbers of reactants/products (*J* ≥ 4) and reactions (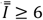), nature of the biochemical network (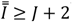) and possible outcomes for each participating reaction (*n* = 3) (**Defs. (3a-3d)**) [36]. A network-specific constraint will depend on the architecture and complexity of the numerical values that comprise the modelled biochemical network. This will include the minimum number of iterations (*u* = *M* ∈ ℕ) needed to unambiguously assign an outcome to every reaction and null space generated subspaces of the stoichiometry number matrix with non-redundant and non-trivial vectors that have been combinatorially summed (**Defs. (4, 5)**; **Figure 1**) [36]. Every row of a subspace is redefined as the *i*^th^-reaction-specific sequence vector, populated and characterized with descriptors (mathematical, statistical) which are then used to select and bin terms to outcome-specific subsets which are labelled as forward (*F*), reverse (*B*) or equivalent (*E*) (**Defs. (6, 7)**; **Figure 1**) [36]. The summed terms of each output-specific subset are then mapped to strictly positive real numbers and constitutes the reaction-specific output vector with a p1-norm that is the probable dissociation constant for the reaction (**Defs. (8, 9)**; **Figure 1**) [36]. “ReDirection” will implement this algorithm iteratively and recursively and compute the probable dissociation constant for every reaction of a user-defined biochemical network (**Figure 1**).

**Figure 1:**
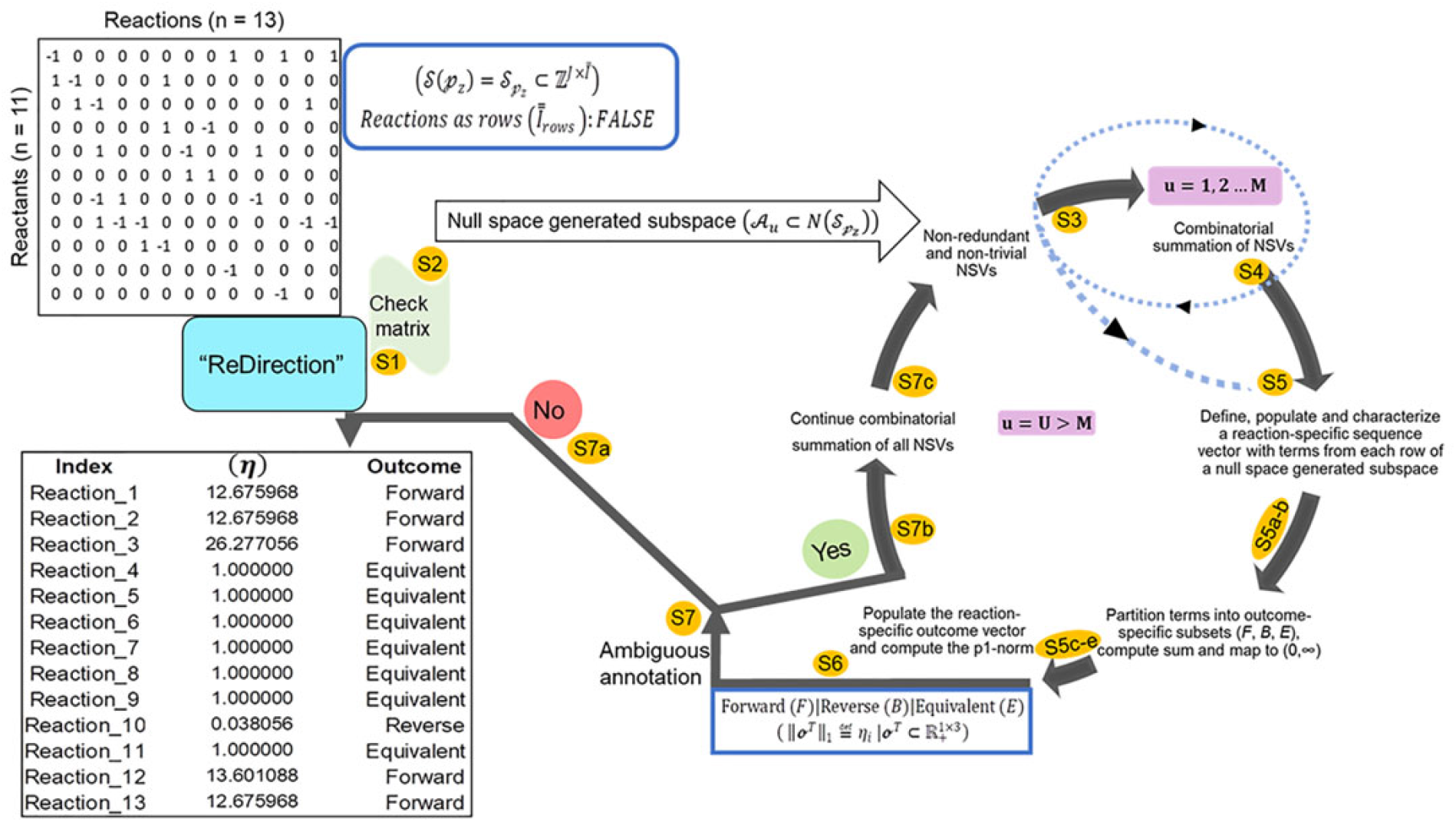
Schematic representation of the steps deployed by “ReDirection” to characterize every reaction of a user-defined biochemical network with the probable dissociation constant. “ReDirection” checks the stoichiometry number matrix that is provided by the user for a modelled biochemical network for compliance with pre-defined constraints. If this is true then “ReDirection” computes a null space generated subspace by excluding all redundant and trivial vectors and combinatorially summing the vectors that remain. “ReDirection” also defines a reaction-specific sequence vector which comprises terms drawn from each row of the resulting subspace. “ReDirection” computes several descriptors (mathematical, statistical) for the numerical values that comprise this vector and partitions these into distinct subsets in accordance with the expected outcomes (forward, reverse, equivalent) for a reaction. “ReDirection” then maps the sum of the terms of each outcome-specific subset to the strictly positive real number and bins these to a reaction-specific outcome vector. The p1-norm of this vector is the probable dissociation constant for a reaction and is used to annotate the same. “ReDirection” accomplishes this recursively and over several iterations until every reaction of the modelled biochemical network has been assigned an unambiguous outcome. **Abbreviations**: η_*i*_, Probable dissociation constant for the *i*^th^-reaction of a user-defined biochemical network; **𝒮**_𝓅_z, User-defined stoichiometry number matrix for a biochemical network; **S1-7**, steps of the algorithm deployed by “ReDirection” to compute the probable dissociation constant and assign outcome to every reaction of a user-defined biochemical network; **NSV**, null space generated subspace vector;

### 2.2 Generic description, availability and guidelines for using ‘ReDirection”

“ReDirection” is freely available and can be updated or installed directly from the graphics user interface (GUI) (R-4.1.x) as “update.packages(‘ReDirection’)” and/or “install.packages(‘ReDirection’)” from any of the CRAN-mirrors. “ReDirection” is built in RStudio (1.4.1717) and tested in R-4.1.x. “ReDirection” comprises three functions (*calculate_reaction_vector, check_matrix, reaction_vector*). The dependencies for “ReDirection” are the packages ‘pracma’, ‘MASS’, ‘stats’ and the *combinations* function from the R-package (‘gtools’). The downloaded package includes detailed documentation of all the functions along with ready-to-use examples and tests of functionality. “ReDirection” will utilize these functions sequentially and process the stoichiometry number matrix of the reactants/products and reactions of a biochemical network that is defined by the user (**Figure 1**). In addition to implementing “ReDirection” locally, several R-scripts will be developed in house to preformat (input, output) and analyse data. Although “ReDirection” is simple to operate, there are a few guidelines that the user needs to be aware of while calculating the probable dissociation constant. “ReDirection” is reaction-centric and requires that the number of reactions of a modelled biochemical network and the number of reactants/products strictly conforms to the pre-defined constraints (**Figure 1**) [36]. Since the user is not expected to validate the stoichiometry number matrix manually, “ReDirection” will undertake this task and do this unequivocally prior to commencing the iterations.

“ReDirection” requires the user to provide a logical argument (TRUE, FALSE) along with stoichiometry number matrix to indicate whether the reactions are to be considered as rows or columns,

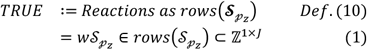

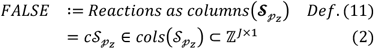

“ReDirection” will utilize this data to assign the appropriate dimensions to the stoichiometry number matrix for a user-defined biochemical network and complete the initial checks of compliance in accordance with the pre-defined constraints (**Step 1**; **Figure 1**) [36]. Another check point, *albeit*, internal is the identification and subsequent exclusion of linear dependent vectors (rows, columns) (**Step 1**; **Figure 1**). These could either be half-reactions (forward, reverse) or an inadvertent introduction/omission by the user (*θ* ∈ ℕ ∪ {0}) (**Def. (13)**). For the case where the user inputs the complete stoichiometry number matrix, i.e., forward- and reverse-reactions,

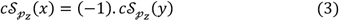

*where,*

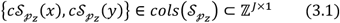

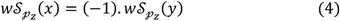

*where,*

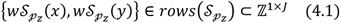

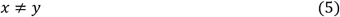

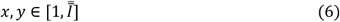

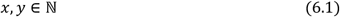

“ReDirection” identifies these and excludes all linear dependent vectors (rows, columns) and rechecks the modified stoichiometry number matrix for constraint compliance (**Steps 1-3**; **Figure 1**). If there are no further deficiencies, “ReDirection” commences the iterations and recursively computes null space generated subspaces by excluding all trivial and redundant vectors and combinatorially summing these (**Steps 3-7**; **Figure 1**),

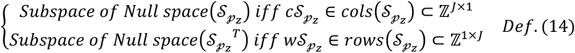

### 2.3 “ReDirection”-based numerical studies on a constrained biochemical network of human Galactose metabolism

Galactose-Glucose interconversion is readily observed within the cell, catalysed by the enzyme UDP-galactose 4-epimerase (*EC* 5.1.3.2) and suggests a biochemical network with several potentially bidirectional reactions [37, 38]. Here, “ReDirection” computes the probable dissociation constants for every reaction of a constrained biochemical network (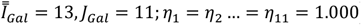) of human Galactose metabolism (**Eqs. (7-9)**). In particular, the effect of perturbing *r*_1_ − *r*_11_ is investigated by introducing the probable reactions *r*_12_ (UDP-Galactose → alpha-D-Galactose 1-phosphate) and *r*_13_ (UDP-Galactose → alpha-D-Galactose 1-phosphate) into the network (**Figures 2a and 2b**). The enzymes (UTP-hexose 1-phosphate uridyltransferase, *EC* 2.7.7.10; UTP-monosaccharide-1-phosphate uridyltransferase, *EC* 2.7.7.64) that mediate the transformation of UDP-Galactose to alpha-D-Galactose 1-phosphate are not significant contributors to human Galactose metabolism. This reaction is mediated by UDP-glucose--hexose-1-phosphate uridyltransferase (*EC* 2.7.7.12) (*r*_9_) and is a major regulatory checkpoint for Galactose-Glucose interconversion (**Figures 2a and 2b**). “ReDirection” computes the probable dissociation constants for every reaction of a constrained biochemical network of human Galactose metabolism (**Figures 2a and 2b**).

**Figure 2:**
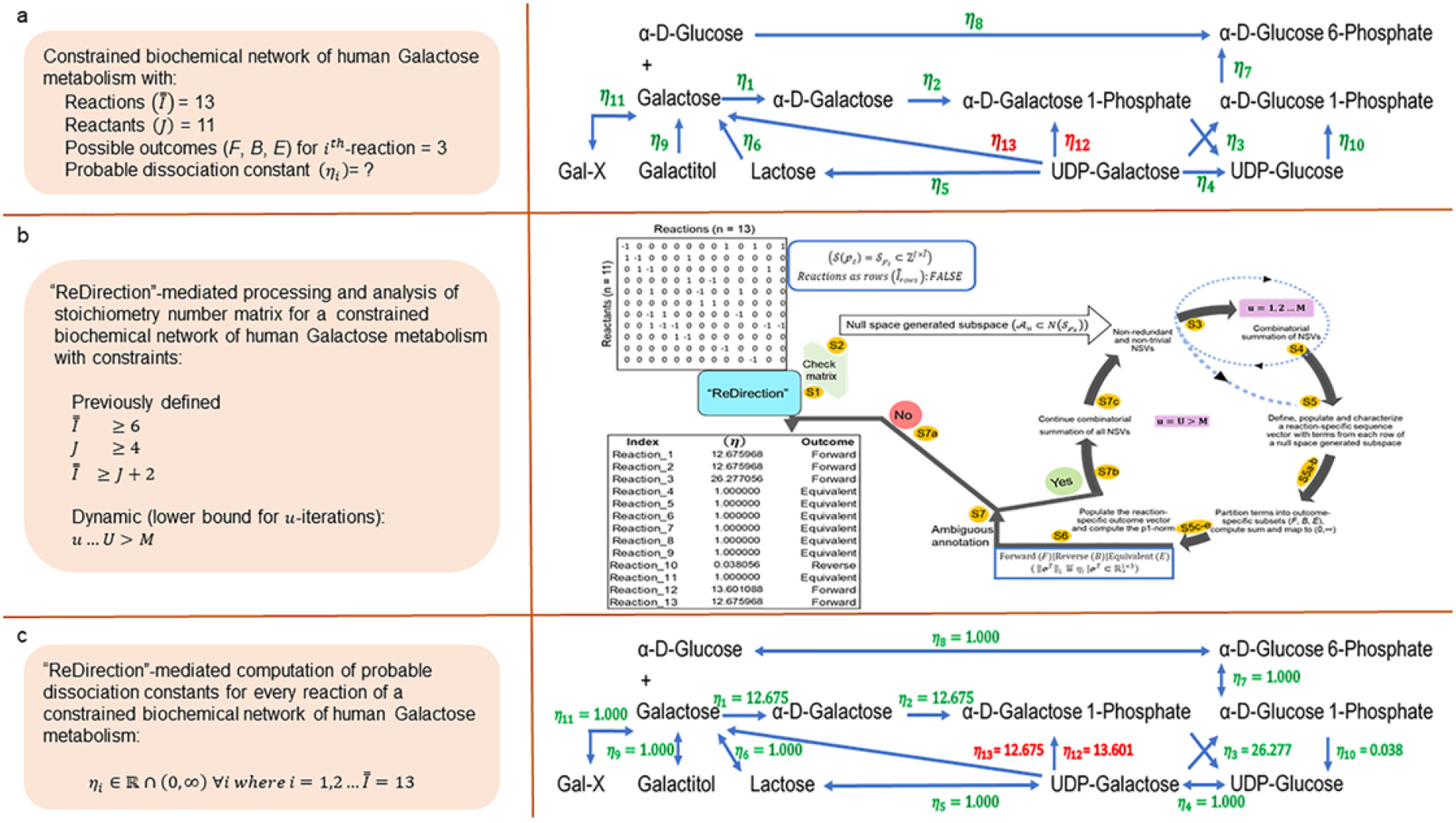
Schematic representation of “ReDirection”-mediated investigation of a constrained biochemical network of human Galactose metabolism. **a)** Definition of a canonical and user-defined constrained biochemical network of human Galacatose metabolism and the matrix of stoichiometric numbers of its reactants/products. The biochemical network for Galactose metabolism in *Homo sapiens* comprises several potentially bidirectional reactions, **b)** The matrix of stoichiometric numbers of the reactants/products of the canonical form of a constrained biochemical network of Galactose in humans is modified and an arbitrarily defined steady-state is investigated by “ReDirection”. In particular, the conversion of UDP-galactose to alpha-D-Galactose 1-phosphate (*r*_12_) and D-Galactose (*r*_13_) via alternate pathways on the unperturbed set of reactions (*r*_1_ − *r*_11_) is investigated and **c)** The output of “ReDirection” is the probable disassociation constant for every reaction of a constrained biochemical network of human Galactose metabolism. These data can be utilized to assign an outcome to every reaction under naive and perturbed (*r*_12_, *r*_13_) conditions. The predictions yield important testable hypotheses in a relatively short time and can be empirically validated. For example, the data suggests that a large proportion of the reactions are equivalent and may therefore, function to regulate Galactose metabolism. Additionally, the net direction observed after computing the probable dissociation constant of every reaction is towards the biosynthesis of UDP-Glucose. **Abbreviations**: **EC**, Enzyme commission; 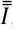, Total number of reactions of a user-defined biochemical network; ***J***, Total number of reactants of a user-defined biochemical network; η_*i*_, Probable dissociation constant for the *i*^th^-reaction of a constrained biochemical network of human Galactose metabolism; **𝒮**_𝓅_z, Stoichiometry number matrix for a constrained biochemical network of human Galactose metabolism; **Gal-X**, Galactose containing di (Galactinol, melibitol, epimelibiose)- or oligo (stachyose)-saccharides which are cleaved by beta-Galactosidase (*EC* 3.2.1.22); **UDP**, Uridine- di-phosphate; **UTP**, Uridine tri-phosphate;

## 3 Results and Discussion

### 3.1 Steps deployed by “ReDirection” to compute the probable dissociation constant for every reaction of a user-defined biochemical network

“ReDirection” will utilize the aforementioned functions sequentially and process the stoichiometry number matrix for the biochemical network that is defined by the user and compute the probable dissociation constant for every reaction (**Figure 1**). This will be done sequentially and is as follows:

**Step 1**: “ReDirection” will check whether the matrix of stoichiometry numbers that the user inputs is compliant with the pre-defined constraints and does not have any linear dependent vectors. If found, “ReDirection” will exclude these. The modified input matrix will be rechecked for constraint compliance.

**Step 2**: “ReDirection” will then compute the null space of the checked/rechecked stoichiometry number matrix of the reactants/products and reactions of the user-defined biochemical network.

**Step 3:** “ReDirection” will process and screen this null space for redundant- and/or trivial-vectors and define a subspace by excluding the same.

**Step 4**: “ReDirection” will combinatorially sum the remaining vectors, i.e., non-redundant and non-trivial, and will repeat **Step 3** for a finite number of *u*-iterations where *u* = *M* ∈ ℕ [36].

**Step 5**: For *u* = *U* > *M* iterations, “ReDirection” will define, populate and compute several descriptors (sum, arithmetic mean, standard deviation) for a reaction-specific sequence vector with terms that are drawn from each row of a null space generated subspace.

> **Step 5a:** “ReDirection” tests each term of an *i*^th^-reaction-specific sequence vector for convergence.
>
> **Step 5b**: If this term diverges and is possesses a numerical value that is greater than 2 standard deviations from the mean, then bin this term into the appropriate outcome-specific (forward/reverse/equivalent) subset.
>
> **Step 5c:** The terms of each outcome-specific subset form a finite series whose sum is computed by “ReDirection”.
>
> **Step 5d**: “ReDirection” then maps these sums to strictly positive real numbers which are then specific for each outcome-specific subset.
>
> **Step 5e**: These outcome-specific numerical measures form the *i*^th^-reaction-specific outcome vector.

**Step 6**: “ReDirection” will compute the p1-norm of the reaction-specific outcome vector and annotate the reaction.

**Step 7:** “ReDirection” will check whether the annotations for all the other reactions of the user-defined biochemical network are unambiguous.

> **Step 7a:** If there is no reaction that has been annotated ambiguously then “ReDirection” will output the predicted outcomes for every reaction of the user-defined biochemical network.
>
> **Step 7b:** If there is a reaction that has been annotated ambiguously then “ReDirection” will continue the iterations.
>
> **Step 7c:** “ReDirection” will combinatorially sum all non-redundant and non-trivial null space generated subspace vectors that remain, define a new subspace and repeat **steps 5-7**.

Although there is no theoretical upper bound on the dimensions of the biochemical network that “ReDirection” can work with, during run-times (i5-10400F processor, clock speed 2.9 GHz, 64-bit, 16 GB RAM) it was observed that when the number of reactions 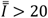 the efficiency, in real-time to unambiguously annotate every reaction of the modelled biochemical network routinely exceeded 2 hours whilst, for lower values of 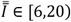 these are completed within 30 minutes (**Eqs. (10, 11)**) [36]. The mathematical rigor underlying the steps used by “ReDirection” to compute the probable dissociation constant is in accordance with the formal algorithm previously described and is presented (**Eqs. (12-27)**; **Table 1**) [36].

**Table 1:** “ReDirection”-based computation of the probable dissociation constant for every reaction of a user-defined biochemical network

|  | Subset (*F*) |  | Subset (*B*) |  | Subset (*E*) |  |
| --- | --- | --- | --- | --- | --- | --- |
| Output-specific subset sum (domain):<br>$x = (.) \in \mathbb{R} \cap (-\infty, \infty)$ | $Case\ 1: \ x_1 > 1$<br>$Case\ 2: \ x_2 > 0$ | (12)<br>(13) | $Case\ 1: \ -x_2 < 0$<br>$Case\ 2: \ x_3 < -1$ | (19)<br>(20) | $\vert x_4\vert \approx 0$ | (24) |
| Linear map:<br>$g: x \in \mathbb{R} \cap (-\infty, \infty) \mapsto y \in \mathbb{R} \cap (0, \infty)$ | $y_F \stackrel{\text{def}}{=} g(x_1 or\ x_2)$<br>$= x_1 or\ x_2$ | (14)<br>(14.1) | $y_B \stackrel{\text{def}}{=} g(-x_2\ or\ x_3)$<br>$= e^{-x_2}\ or\ e^{x_3}$ | (21)<br>(21.1) | $y_E \stackrel{\text{def}}{=} g(x_4)$<br>$= e^{x_4}$ | (25)<br>(25.1) |
| Range:<br>$y(.) \in \mathbb{R} \cap (0, \infty)$ | $y_F \in \mathbb{R} \cap (1, \infty)$ | (15) | $y_B \in \mathbb{R} \cap (0, 1)$ | (22) | $y_E \in \mathbb{R} \cap \{1\}$ | (26) |
| Reaction-specific outcome vector:<br>$\boldsymbol{o}^T$ | $[g(x_1 or\ x_2)\ g(-x_2 or\ x_3)\ g(x_4)]^T = [(x_1\ or\ x_2)\ (e^{-x_2}\ or\ e^{x_3})\ e^{x_4}]^T \quad (16)$<br>$= [y_F\ y_B\ y_E]^T \quad (16.1)$ | | | | | |
| Prediction | Forward |  | Reverse |  | Equivalent |  |
| Probable dissociation constant (p1-norm):<br>$\ \boldsymbol{o}^T\ _1 = \eta_i \in \mathbb{R} \cap (0, \infty)$ | $Case\ 1: \ x_1 + 0 + 0$<br>$= \eta_i \in \mathbb{R} \cap (1, \infty)$<br>$Case\ 2: \ x_2 + e^{-x_2} + 0$<br>$= \eta_i \in \mathbb{R} \cap (1, \infty)$ | (17)<br>(17.1)<br>(18)<br>(18.1) | $0 + e^{x_3} + 0$<br>$= \eta_i \in \mathbb{R} \cap (0, 1)$ | (23)<br>(23.1) | $0 + 0 + e^{x_4}$<br>$= \eta_i \in \mathbb{R} \cap \{1\}$ | (27)<br>(27.1) |

**Abbreviations:**
- $x(.)$ : Real-valued numeral of a null space generated subspace - $g: x(.)$ : Linear map for real-valued numeral of a null space generated subspace - $y(.)$ : Strictly positive mapped real-valued numeral - $\sigma^T$ : Reaction-specific outcome vector - $i$ : $i^{th}$ -reaction of a user-defined biochemical network - $\eta_i$ : Probable dissociation constant for the $i^{th}$ -reaction of a user-defined biochemical network - $F, B, E$ : Outcome-specific subsets (forward, $F$ ; reverse, $B$ ; equivalent, $E$ )

### 3.2 “ReDirection”-based characterization of a constrained biochemical network of human Galactose metabolism

The probable dissociation constants that are computed for a constrained biochemical network of human Galactose metabolism by “ReDirection” offer several interesting insights into the crosstalk between Glucose and Galactose (**Figure 2c**). There is a significantly larger fraction (≈ 63%; *I*_cat_ = 7) of equivalent reactions as compared to the forward (≈ 27%; *I*_cat_ = 3)- and reverse (≈ 10%; *I*_cat_ = 1)-reactions (**Eqs. (28-30); Figure 2c**). An intriguing observation from these data is the directional preference of the forward (η_1_ = η_2_ ≈ 12.7, η_3_ ≈ 26) and reverse (η_10_ ≈ 0.04)-reactions towards the synthesis of UDP-Glucose (**Eqs. (31, 32); Figure 2c**). This, when coupled to the equivalent and sequential conversions to UDP-Galactose, Lactose and Galactose (η_4_ = η_s_ = η_6_ ≈ 1.000) will ensure that there is minimal change to the pool of Galactose-containing complex-carbohydrates and -lipids (glycosphingolipids, gangliosides, cerebrosides, mucopolysaccharides) (**Eq. (33)**). Additionally, since the magnitude of the probable dissociation constant for η_3_ is twice that of η_1_ and η_2_ 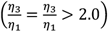 the conversion or utilization of alpha-Galactose 1-phosphate is faster that its synthesis (**Eqs. (34, 35)**). This will, by the law of mass action also ensure that the net flux of the network is towards the biosynthesis of UDP-Glucose (**Figure 2c**).

The atypical reactions (*r*_12_, *r*_13_) via the computed probable dissociation constants (η_12_, η_13_) will function to perturb the constrained biochemical network of human Galactose metabolism (**Figure 2**). The computed data suggests that the preferred outcome for these reactions is “forward” and towards the synthesis of Galactose 1-phoshate (η_12_ ≈ 13.6) or Galactose (η_13_ ≈ 12.7) from UDP-Galactose (**Eqs. (36, 37)**; **Figure 2c**). Clearly these reactions can either complement or compensate, where applicable, the reactions *r*_1_ and *r*_2_ (η_1_ = η_2_ ≈ 12.7) (**Eqs. (38, 39)**; **Figure 2c**). This predilection towards synthesizing Galactose may, along with the activity of UDP-galactose 4-epimerase, constitute a plausible explanation for the milder clinical profile of the inborn errors of metabolism for Galactose biosynthesis and utilization [42, 43]. Beta-Galactosidase (*EC* 3.2.1.23), on the other hand, prefers to cleave Lactose to generate D-Galactose (*r*_6_) rather than homopolymeric galactans (*r*_13_) (**Figure 2a**).

### 3.3 The probable dissociation constants for a biochemical network are suitable indices of biochemical function

The probable dissociation constants for a modelled biochemical network are useful indices of net directionality and reaction-rate [6-10, 36]. A potentially novel application for these data is the conduct of simulation studies with the stochastic simulation algorithms to compute the trajectories of the reactants/products [39-41]. This approach has yielded interesting insights into the export of high-affinity peptides to the plasma membrane by the major histocompatibility complex-I (MHC1) and the directed chemotaxis of a phagocyte towards a noxious stimulus [40, 41]. However, these studies mandate by definition, the use of every possible reaction which will preclude direct usage of the “ReDirection”-modified stoichiometry number matrix since,

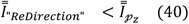

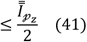

If we rewrite the constraints as defined by the algorithm in terms of this reduced set of reactions,

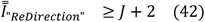

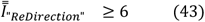

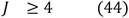

These modifications, if needed, will imply that the set of reactions for a user-defined biochemical network 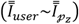 is,

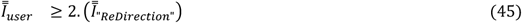

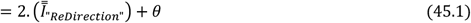

*where,*

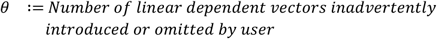

Clearly, the user-defined set of reactions for a modelled biochemical network can be used to simulate the modelled biochemical network after including the excluded half-reactions whilst excluding other redundant vectors,

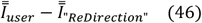

These will have to be included explicitly [39-41]. In other words,

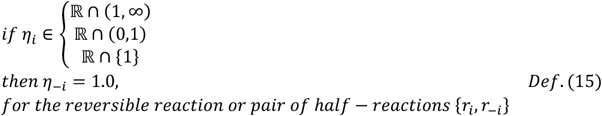

The lower bound or minimum number of iterations (*u* = *M*) that “ReDirection” must conduct before concluding the calculations is determined when the least upper (> 1)- or greatest lower (< -1)-bounds for each reaction-specific sequence vector holds unequivocally or the number of positive- and negative-terms are almost equal for *u* = *U* > *M* (**Def. (16)**) [36]. This is dependent on the complexity of the null space generated subspace and therefore, indirectly on the user-defined stoichiometry number matrix for the modelled biochemical network.

## 4 Conclusions

“ReDirection” is a numerically robust R-package that will compute the probable disassociation constant directly from the null space generated subspace of a stoichiometry number matrix of a user-defined biochemical network and assign an outcome to every reaction. Whilst mathematical rigor is ensured at all steps, biological relevance is maintained by utilizing parameters and metrics in accordance with established kinetic paradigms. “ReDirection” computes the probable dissociation constant from first principles and can be used to compare biochemical networks under varying intracellular environments (naive, perturbed), between cells and across taxa. Although computationally intense and possibly intractable for larger networks, the predictions are reasonably rapid for fewer reactions and are completed quickly in a desktop environment. Future investigations will strive to improve upon the computational time, investigate perturbations, and validate some of the findings by simulation studies. “ReDirection” is not discovery-based and is better suited to addressing known and often empirically intractable biochemical problems *in silico* with simulations or generating testable hypotheses in a laboratory setting.

## Author(s) contributions

### Siddhartha Kundu (SK)

Conceptualization, Methodology, Software, Resources, Formal analysis, Data curation, Validation, Visualization, Investigation, Writing-Original draft preparation, Writing-Reviewing and Editing, Funding acquisition.

## Acknowledgement(s)

This study was supported by an extramural grant from the Science and Engineering Research Board (SERB), Department of Science and Technology, Government of India under the Mathematical Research Impact-Centric Support (MATRICS) scheme to SK (MTR/2021/000290).

